# Comparative analysis of protein evolution and RNA structural changes in the genome of pre-epidemic and epidemic Zika virus

**DOI:** 10.1101/050278

**Authors:** Arunachalam Ramaiah, Lei Dai, Deisy Contreras, Sanjeev Sinha, Ren Sun, Vaithilingaraja Arumugaswami

**Affiliations:** Centre for Infectious Disease Research, Indian Institute of Science, Bangalore 560 012, India.; Department of Molecular and Medical Pharmacology, David Geffen School of Medicine, University of California at Los Angeles, CA 90095.; Department of Ecology and Evolutionary Biology, University of California at Los Angeles, CA 90095; Board of Governors Regenerative Medicine Institute, Cedars-Sinai Medical Center, Los Angeles, CA 90048.; All India Institute of Medical Sciences, New Delhi, India.; Department of Surgery, Cedars-Sinai Medical Center, Los Angeles, CA 90048.; Department of Surgery, David Geffen School of Medicine, University of California at Los Angeles, CA 90095.; Both authors contributed equally to this work.

## Abstract

Zika virus (ZIKV) infection is associated with microcephaly, neurological disorders and poor pregnancy outcome^1-3^ and no vaccine is available. Although ZIKV was first discovered in 1947, the exact mechanism of virus replication and pathogenesis still remains unknown. Recent outbreaks of Zika virus in the Americas clearly suggest a better adaptation of viral strains to human host. Understanding the conserved and adaptive features in the evolution of ZIKV genome will reveal the molecular mechanism of virus replication and host adaptation. Here, we show comprehensive analysis of protein evolution and changes in RNA secondary structures of ZIKV strains including the current 2015-16 outbreak. To identify the constraints on ZIKV evolution, selection pressure at individual codons, immune epitopes, co-evolving sites, and RNA structures were analyzed. The proteome of current 2015/16 epidemic ZIKV strains of Asian genotype is found to be genetically conserved due to genome-wide negative selection on codons, with limited positive selection. Predicted RNA structures at the 5’ and 3’ ends of ZIKV strains reveal substantial changes such as an additional stem loop which makes it similar to that of Yellow Fever Virus. Concisely, the targeted changes at both the amino acid and the RNA levels contribute to the better adaptation of ZIKV strains to human host with an enhanced neurotropism.

## MAIN TEXT

Zika virus (ZIKV) is an emerging disease strongly linked to poor pregnancy outcomes and birth defects specifically microcephaly, a condition with an abnormally small head^1,4^. The virus is transmitted by mosquito vector *Aedes aegypti* and sexual contact^5-7^. Zika viral particle contains a positive sense, single-stranded RNA genome of about 10.7 kb^8^. The genome is organized as 5’UTR-C-prM-E-NS1-NS2A-NS2B-NS3-NS4A-2K-NS4B-NS5-3’UTR, with untranslated regions (UTR) flanking a protein-coding region. The latter encodes a single polyprotein (3423 amino acids) that is co-and post-translationally cleaved by cellular and viral proteases into multiple peptides. 5’ and 3’UTR stem loop RNA structures have been shown to be critical for the initiation of viral genome translation and replication in other Flaviviruses.

ZIKV was originally isolated from a sentinel rhesus monkey in the Zika forest of Uganda in 1947 and the first human case based on serological evidence was reported in 1952^9^. Detailed history of ZIKV outbreaks and phylogeny are described ^10^ ^11,12^. Currently, the ZIKV strains are classified into three major genotypes or lineages, including West African (Nigerian cluster), East African (MR766 prototype cluster), and Asian ^11^. For the first time in 2007, ZIKV outbreak was reported in the Pacific region at Yap Island in the Federated States of Micronesia^13^. Subsequent epidemics have occurred in French Polynesia in 2013 and the Americas including Brazil, Columbia, Guatemala and Puerto Rico in 2015^7,14^. Genetic analysis revealed that the Asian genotype of ZIKV is responsible for the Pacific outbreaks^11,12,14-17^.

The Zika virus African lineage (strain MR766) shares 96% and 56% amino acid identity with that of the ZIKV Asian lineage and dengue virus 2, respectively. Given the high mutation rate of RNA viruses^18^, this suggests that highly conserved features of the ZIKV genome may reflect elements critical for virus replication. In contrast, genetic variations in the ZIKV genome may reveal how viruses have adapted to new environments, such as human hosts. Sylvatic cycle comprising non-human primates and *Aedes* mosquitos with occasional involvement of human host was the original mode of transmission. Subsequent molecular adaptation of ZIKV to human host may be responsible for increase in virulence and current outbreaks. Previous studies have shown patterns of selection pressure, recombination events, and glycosylation in pre-epidemic ZIKV strains sampled from 1947-2007^12^. Recent studies on the epidemic strains have mainly focused on the phylogeography of ZIKV isolates^11^, ^15,19^. However, detailed analysis on the molecular evolution of epidemic ZIKV strains is still lacking^17^.

In this study, we performed comprehensive analysis on protein evolution and changes in RNA secondary structures of 46 ZIKV strains isolated from 1947 to 2016, including 20 strains responsible for the current 2015-16 outbreak. We demonstrate how evolutionary forces and constraints have shaped the genome of current epidemic ZIKV strains, including genome-wide negative selection on amino acid substitutions, co-evolving sites in the proteome, immune epitopes, as well as the roles of RNA structures in host adaptation and virus replication. We further discuss the implications of our results in the broader context of vaccine design and emergence of RNA viruses.

Identifying the origin and distribution of 2015/16 epidemic ZIKV strains would be crucial for diagnostics, vaccine development and disease management^12^. The evolutionary relationship of 46 ZIKV strains isolated from 1947 to 2016 showed one clade was formed by 28 Asian strains which included all twenty 2015/16 epidemic ZIKV strains, while the other by the 18 African strains (Fig. 1 and Supplementary Fig. 1). Sequence alignments also confirmed deletions in the envelope protein glycosylation motif of the African lineages, which is not the case for Asian lineage (Supplementary Fig. 2). The polyprotein synthesized by the coding region of 2015/16 ZIKVs strains had 99.6%-99.9% similarities, which denoted genetic conservation among current viral strains. These circulating strains had close genetic associations with non-epidemic human H/PF/2013 ZIKV (KJ776791) and differed in only 16 non-synonymous substitutions. Our findings also indicate that the Asian lineage has dispersed to several countries across the Asian and the American continents, whereas the dissemination of the African lineage was restricted to the countries within the African continent.

**Figure 1.**
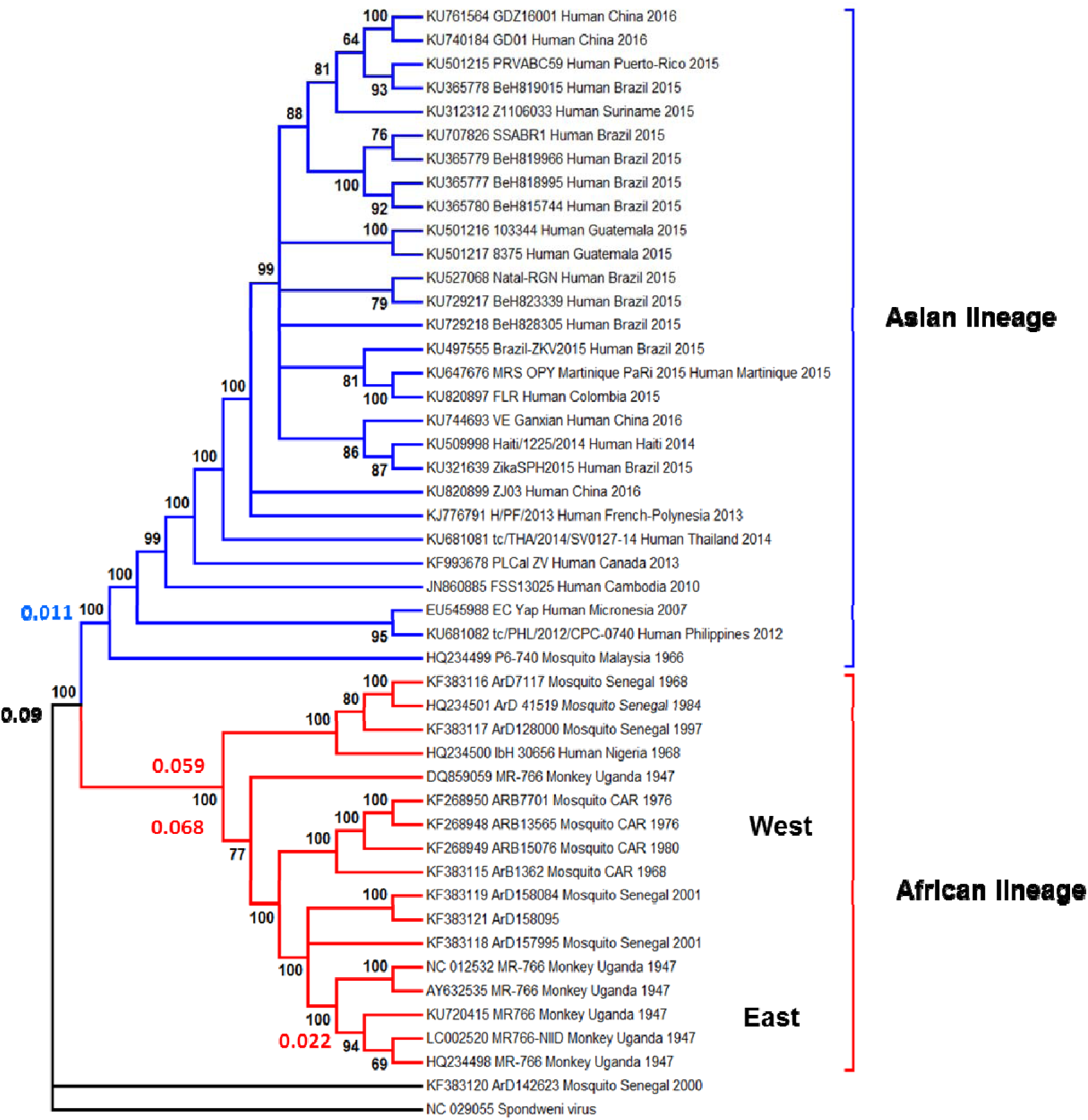
Phylogenetic relationships of Zika viral strains of East African, West African and Asian lineages. Bootstrap consensus NJ tree was reconstructed with 1000 bootstrap replications, with support values >60% shown in the nodes. The African and Asian clades consist of 18 and 28 strains that were colored in red and blue, respectively. The Spondweni virus (NC_029055) was used as an out group. A total of 20 ZIKV responsible for the current 2015/16 outbreak were clustered together within Asian clade. Whereas, the West African strains were segregated from East African strains. The calculated average of genetic distance within the strains from three clades and an overall mean were shown near the clades.

We evaluated genome-wide amino acid substitutions to understand the nature of selection pressure acting on each codon. Higher rate of synonymous (dS) substitutions than non-synonymous (dN) substitutions is usually considered as the signature of negative selection, also known as purifying selection. For the 46 ZIKV genomes isolated from 1947 to 2016, 70% of the 3423 codons were found to be polymorphic, indicating substantial evolution in the viral proteome (Fig. 2; Supplementary Table 1). Among the three different lineages, we found that 46-115 codons (1.3%-3.4%) were identified with higher rate of non-synonymous substitutions (dN>dS), while 560-1664 codons (16.4%-48.6%) had higher rate of synonymous substitutions (dN<dS).

**Figure 2.**
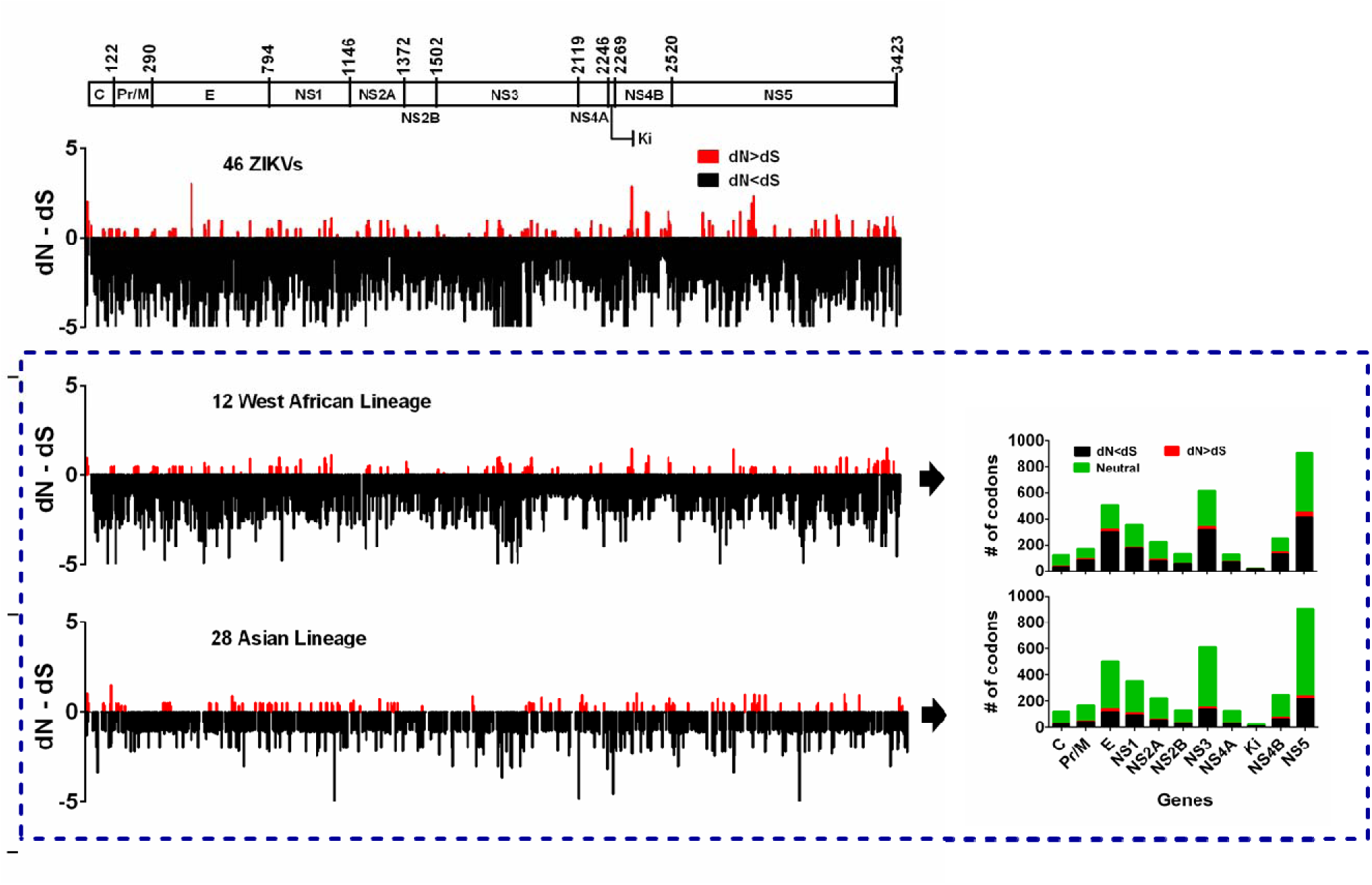
Schematic representation of dN-dS test statistics for 3423 codons of Zika viral strains. The dN-dS value for each of 3423 codons indicated as a bar either above or below the X-axis. The positive values indicate an excess of non-synonymous substitutions (red bars) in the codons, whereas, negative values below the horizontal line indicate an excess of synonymous substitutions (black bars). Bars were arbitrarily trimmed at 5 to save space. A total of 46 strains including 12 West African and 28 Asian viruses were shown in the graph. A schematic of ZIKV genome is aligned on the top of the graphs. dN-dS scores are presented in Y-axis. The stacked bar diagrams (right side) show the number of codons with dN<dS (black), dN>dS (red) and neutral (green) selected codons in each gene of Zika viral strains belong to West African and Asian lineages.

We carried out statistical tests of dN-dS for codons from each of the 11 protein segments of the Asian and West African lineages (Supplementary Text 1). For example, in the Asian lineage, 31 (26%) of 122 codons in the capsid (C) were polymorphic. Among the polymorphic codons, a higher rate of non-synonymous (dN>dS) and synonymous (dN<dS) substitutions was observed in 7 (6%) and 24 (20%) codons, respectively (Fig. 2, Supplementary Fig. 3a-d). The selection profiles for all the other segments in the Asian lineage were found to be qualitatively similar. We also observed an overall similar pattern of dN and dS in the West African lineage.

The nature of selection pressure exerted on each codon of different ZIKV data sets can be measured by obtaining the ratio of dN and dS (Supplementary Table 2a). This analysis showed that the dN/dS ratios of all data sets, especially both Asian and West African lineages were much smaller than one, with the highest values observed in the capsid (0.239) of the Asian lineage and in the pre-membrane (0.118) of West African lineages (Supplementary Table 2b). Our results suggest that non-synonymous mutations have a lower probability of being fixed than the synonymous mutations, as changes in the protein sequence are on average more likely to decrease the replicative fitness of ZIKV. Thus, the proteome of the epidemic ZIKV strains is genetically conserved due to genome-wide negative selection on amino acid substitutions.

We subsequently examined the potential selection pressure applied by immune system on viral epitopes. We identified 106 CD4 T cell epitopes (TCEs) and 5 CD8 TCEs that had higher probability to induce an immune response (Supplementary Text 2). Among the 106 highly immunogenic CD4 TCEs, 10 peptides contained negatively selected amino acid sites. This observation indicates that immune escape variants can be strongly selected against due to fitness cost.

To identify higher-order constraints on the evolution of ZIKV proteome, we inferred potentially coevolving amino acid sites in the African and Asian lineages by computing correlation between the variations at each pair of codons (Fig. 3a,b). The effect of phylogeny on site linkage was cleaned by removing the contribution of the largest eigenvalue(s) from the correlation matrix (Supplementary Figs. 4 and 5). Positive correlation between two sites implies that the corresponding double mutants are observed more frequently than if the mutations were to occur independently. In contrast, negative correlation between two sites implies that the double mutants are observed less frequently than if the individual mutations were to arise independently. In both cases, the dependence of one site on another implies epistasis, i.e. the fitness effect of multiple mutations is considerably different from the additive effect of single mutations.

**Figure 3.**
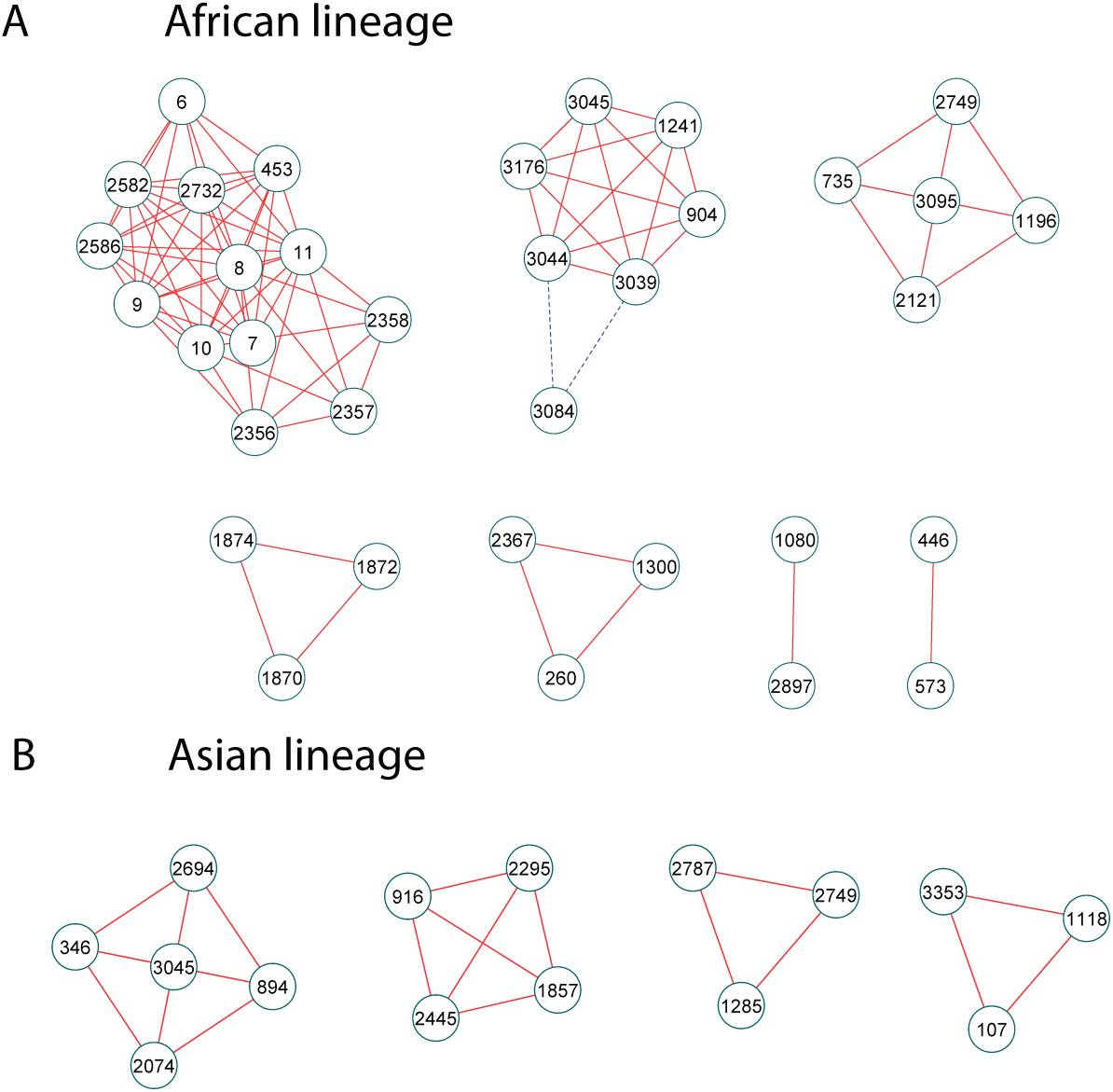
Network of coevolving sites of ZIKV polyprotein. (a) African lineage. (b) Asian lineage. The nodes are labelled by site number of ZIKA polyprotein (from 1 to 3423). The edges indicate significant positive correlation (red solid lines) or negative correlation (blue dashed lines) between two sites.

Among the 80 polymorphic sites in the African lineage, seven groups of sites showed significant pairwise correlation. The observed correlation between physical proximal sites, such as codon 1870-1872-1874 (Fig. 3a), may reflect epistatic interactions within a protein, NS3 in this case. Correlation between sites of different proteins may suggest potential protein-protein interactions, which can be further studied via mutagenesis and protein structures.

There are 32 polymorphic sites in the Asian lineage due to the limited genetic diversity (Fig. 3b). Given the very different genetic background of African and Asian lineages, it is not surprising to see that the two coevolution networks are composed of different sites. The similarity between the two networks is that most of the pairwise interactions are positive. One well-known example of positive interactions is compensatory mutations, where the fixation of a second mutation rescues the preceding deleterious mutation^20^. Interpreting coevolving site data requires additional caveats which are discussed in Supplementary Text 3. The observed correlation can be further improved by utilizing larger sample size of fully sequenced Zika viral genomes.

Subsequently, we examined the RNA structures. Using the full-genome sequence of ZIKV (Supplementary Table 3), we performed a comparative analysis of RNA secondary structures in the 5’ and 3’ UTR (Fig. 4). The RNA structure was analyzed for each strain by IPknot, one of the most accurate single-sequence prediction method^21,22^. The dendrogram is constructed based on the dissimilarity between RNA secondary structures. Similar to the phylogenetic tree based on genetic distance of protein sequences, the RNA structures of the African lineage and Asian lineage are separated into two clusters, suggesting that the RNA structures in UTR have evolved in parallel to the proteome.

**Figure 4.**
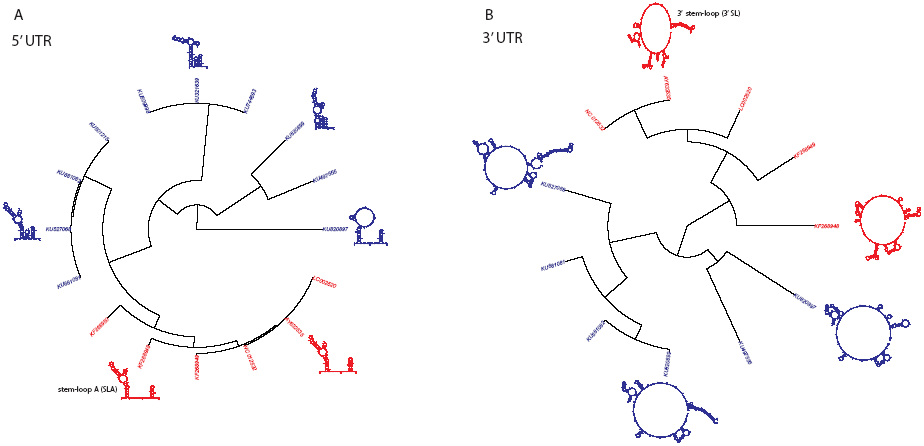
Comparison of RNA secondary structures in 5’ and 3’ UTR. The RNA secondary structure in (a) 5’ UTR or (b) 3’ UTR is predicted by IPknot based on a single sequence. The fan dendrogram shown is based on the distance matrix among predicted structures given by RNA distance in Vienna RNA package. Red: strains of African lineage. Blue: strains of Asian lineage.

Stem Loop A (SLA) in the 5’ UTR is conserved among African and Asian strains, which is also observed in other Flavivirus genomes (Fig. 4a). Compared to the African strains, there is an additional stem loop in the Asian strains before the coding region. Thus, the 5’ UTR structure of the epidemic strains seems to have evolved and become more similar to Yellow Fever Virus^23^. For 3’ UTR, a long stem loop at the 3’ end of the genome (3’ SL) is conserved among all strains (Fig. 4b). However, the sequence and the RNA structure in other parts of 3’ UTR are much less conserved and include complex pseudoknots, in accordance with the general pattern observed in other Flavivrus^24^. The 3’ UTR has been proposed to play a critical role in host adaptation and pathogenesis of Flavivirus^25-27^, and its huge variations may indicate conflicting selection in different hosts. We found that RNA structures of strains isolated in three different hosts, including Sentinel monkey (African lineage), mosquito (African lineage) and human (Asian lineage), displayed many variations except 3’ SL (Supplementary Fig. 6). Although the mechanism underlying these changes are largely unknown, one possible explanation provided by studies in Dengue virus points to subgenomic viral RNAs produced near the 3’ end, where the cellular exoribonuclease is stalled by pseudoknots in 3’ UTR^28^.

In additional to SLA and 3’ SL, we found that RNA structural elements involved in genome cyclization are highly conserved across African and Asian strains of ZIKV (Supplementary Fig. 7). Base-pairing of complementary sequences from the 5’ and 3’ terminals results in RNA cyclization, which is essential for RNA replication of Flavivirus (Villordo and Gamarnik 2009). The two pairs of complementary sequences, including 5’-3’ complementary sequences (5’-3’ CS) and 5’-3’ upstream AUG region (5’-3’ UAR)^23^, are identical in all the strains we examined.

We further extended the analysis of RNA structures to the coding region (Supplementary Fig. 8). From the predicted RNA structure of 300 bp at the 5’ end of coding region (i.e. capsid protein) or at the 3’ end of coding region (i.e. NS5 protein), we identified a few conserved elements by comparing the consensus structure of African strains with Asian strains. The capsid hairpin downstream of the start codon, which is known to be involved in translation initiation, is conserved^23^. In addition, we identified 3 other conserved hairpins in the capsid protein and two conserved hairpins in the NS5 region. The prevalence of conserved structures in coding regions is consistent with previous findings in other RNA viruses, including HCV and HIV^29^. Whether these conserved structures of ZIKV are functional in virus replication remain to be elucidated.

Experiments to probe the function of RNA structures in ZIKA genome of various lineages in different hosts will elucidate the importance of their roles in virus replication (e.g. cyclization), host adaptation (e.g. pseudoknots in 3’ UTR) and emergence. In future studies, the prediction of RNA secondary structures can be greatly improved by additional experimental constraints, such as SHAPE^30^. Our detailed bioinformatics approach provides a framework for comparing and sub-classifying clinical isolates based on RNA structures and developing sub-unit vaccine candidates.

Our comprehensive analysis of various Zika virus isolates identifies specific amino acid residues and non-coding regions that can contribute to the observed changes in virulence and host tropism. Reverse genetics approach by engineering point mutations and inter and intra-genotype chimeric viruses would further address the biological significance of these observed changes. Deep sequencing analysis of the viral quasi-species present in various human organs including fetal brain, placenta, blood and bodily fluids can further improve our understanding of: 1) intra-host genetic diversity of virus; 2) the relation between intra-host genetic diversity and pathogenicity; 3) transmission bottleneck.

## ACKNOWLEDGMENTS

This work was funded by the Cedars-Sinai Medical Center’s Institutional Research Award to V.A., and NIH PO1 CA177322 to R.S. L.D. was supported by HHMI Postdoctoral Fellowship from Jane Coffin Childs Memorial Fund for Medical Research.

## MATERIALS AND METHODS

**Phylogenetic analysis of Zika viral strains**. A total of 46 existing ZIKV genomic sequences including 20 ZIKVs (2015/16 isolates) that are responsible for the current epidemic outbreak were retrieved from GenBank (www.ncbi.nlm.nih.gov/genbank) (dated 10-Mar-2016). The considered ZIKV strains were isolated from human, monkey and different species of mosquito from 1947 to 2016. Additionally, genome sequence of single Spondweni virus (NC_029055) from the Flavivirus family was included in our study as an out group species. The strain name, accession number, host, year of isolation, country and strain lineage information listed in the Supplementary Table 3. Multiple sequence alignments for 47 viral genomes were performed using MUSCLE^1^, subsequently the phylogenetic trees were reconstructed using Neighbor Joining (NJ), Maximum Parsimony (MP) and Maximum Likelihood (ML) approaches with 1000 bootstrap supports. The NJ tree was reconstructed under Tajima-Nei model^2^. GTR+G+I were identified as the best model using Model Test program for reconstructing the ML tree using MEGA6^3^. The genetic distance (GD) between the strains in each clade and between the clades (East African, West African and Asian lineages) were calculated in MEGA6 using the NJ algorithm under Tajima-Nei model^2,3^.

**Estimating dN-dS score and dN/dS ratio**. The ML computations of dN and dS were conducted using HyPhy software package^4^ implemented in MEGA6^3^. The dN-dS score and dN/dS ratio were estimated for detecting ZIKV codons that have undergone positive or negative selection, here dS is the number of synonymous substitutions per site (s/S) and dN is the number of non-synonymous substitutions per site (n/N).

**Detection of site-specific selection pressure on epidemic human Zika viral genomes**. The selection pressure operating on each codon/amino acid site of epidemic 2015/16 ZIKVs was detected by computing the dN/dS (ω) ratio. The following multiple statistical methods from Datamonkey^5^ were used for the selection analyses: SLAC, FEL^6^, IFEL^7^, FUBAR^8^, MEME^9^. These methods are described in detail elsewhere^6-12^.

**Analysis of co-evolving sites**. Amino acid sequences of coding region (from site 1 to 3294) are aligned by MUSCLE. The sequences are separated into the African lineage (17 sequences) and the Asian lineage (28 sequences) to preserve phylogenetic homogeneity. The gaps are filled by amino acid of the consensus sequence of each lineage. The alignment is then transformed into a matrix of binary variables. For each site, if the amino acid is the same as the consensus sequence, it is represented by 0; otherwise, it is represented by 1. The analysis of correlation matrix is restricted to polymorphic sites, which we define as positions at which at least 2 sequences have non-consensus amino acid.

The effect of phylogeny to the correlation matrix is cleaned up by removing the contributions of the largest two eigenvalues (African lineage) or the largest eigenvalue (Asian lineage). The cleaned correlation matrix is reconstructed by the remaining eigenvalues. To compare the uncleaned and cleaned correlation matrix (Supplementary Figs. 4 and 5), the sites are assigned into sectors by the loadings of eigenvectors. The correlation between two sites is considered significant if the following criterions are met: 1) in the cleaned correlation matrix, the magnitude of the correlation is among the top 5%; 2) in the un-cleaned correlation matrix, the magnitude of correlation is larger than 0.5 to ensure that Z score is larger than 2. The Z score for the correlation between two sites is estimated by 1000 random permutations at each column of the sequence alignment. The analyses are performed by custom MATLAB scripts. The network of sites with significant correlation is visualized by Cytoscape^13^.

**RNA secondary structures**. Sequences of 5’ UTR (at least 100bp) or 3’ UTR (at least 400bp) are used to predict the RNA secondary structure by IPknot^14^ under default settings. Concatenated sequences of 5’ terminal (170 bp) and 3’ (130 bp) are used to predict the consensus RNA structure after cyclization. When pseudoknots are allowed, the RNA structure is predicted with nested pseudoknot. The RNA structure is visualized by VARNA^15^. The distance matrix between RNA secondary structures is calculated by RNAdistance in ViennaRNA package 2.0^16^. The dendrogram based on the distance matrix is generated by APE package in R^17^.

**Predictions of CD4 and CD8 T cell epitopes**. We have considered 6 proteins/segments: C, E (domain), NS2A, NS3-serine protease, NS4A, and NS5 from an epidemic human BeH819966 ZIKV (KU365779), as a representative of genetically conserved all 20 human 2015/16 ZIKVs to predict all possible CD8 T cell epitopes (TCEs) and CD4 TCEs using MHC-I and MHC-II TCE prediction tools ^18^ under default conditions implemented in Immune Epitope Database (IEDB)^19^.

The predicted peptides were tested for their immunogenicity using immunogenicity prediction tool^20^, here the peptide with higher immunogenicity score indicates a greater probability of mounting an immune response.

## SUPPLEMENTARY FIGURES

**Supplementary Figure 1. ML and MP phylogenetic trees of Zika viral strains**. The ML tree was reconstructed under best GTR+G+I model. In both ML and MP trees, the strains were segregated into two major genotypes, African and Asian. No significant differences found in the topology of NJ, ML and MP trees.

**Supplementary Figure 2. Deletion of a glycosylation motif in E protein**. Sequence alignment of E protein (amino acid positions 140-191) of various strains has shown 4-6 codons/amino acids deletion in the glycosylation site of African lineage. Dashes indicate deleted residues.

**Supplementary Figure 3. Schematic representation of dN-dS test statistics for Zika viral strains (African lineage)**. The dN-dS value for each of 3423 codons indicated as a bar either above or below the X-axis. (a) Autonomous deep view of dN-dS test statistic graphs for codons from each of 11 genomic segments (C, Pr/M, E, NS1, NS2A, NS2B, NS3, NS4A, Ki, NS4B, NS5) of 28 Asian ZIKVs genomes. Codons with excess of dN or dS are shown along with neutrally evolved codons (zero as a score). The X-axis shows the respective number of codons and their range in the genes/segments. (b) 18 African lineage strains (including both East and West African lineages) and 6 East African lineage strains are shown in the graph. Y-axis presents the values of dN-dS scores. The Venn diagrams show the comparison results of overabundance of (c) non-synonymous and (d) synonymous substitutions in the codons from phylogenetically related Asian and West African lineages.

**Supplementary Figure 4. Removing the effect of phylogeny from the correlation matrix (African lineage)**. (a) Multiple sequence alignment (MSA) of polymorphic sites. The analysis of correlation matrix is restricted to polymorphic sites, which we define as positions at which at least 2 sequences have non-consensus amino acid. (b) Correlation matrix before cleaning up phylogeny. (c) Eigenvalue spectrum. The red line indicates the highest eigenvalue observed in the correlation matrix of 1000 randomized MSA, where the columns are shuffled. (d) Correlation matrix after cleaning up phylogeny. To visualize the contribution from the largest eigenvalue(s), the sites are re-ordered according to the loadings of the top 4 eigenvectors.

**Supplementary Figure 5. Removing the effect of phylogeny from the correlation matrix(Asian lineage)**. The notations are the same as in Supplementary Figure 4.

**Supplementary Figure 6. RNA secondary structure in 5’ and 3’ UTR of strains isolated from different hosts**. RNA secondary structures of three strains isolated from (a) Sentinel monkey, (b) mosquito and (c) human are shown. Prediction of RNA structures in 3’ UTR is allowed to have nested pseudoknots.

**Supplementary Figure 7. Cyclization of ZIKV genome**. (a) Predicted secondary structure of 5’ terminal (170 base pairs) and 3’ terminal (130 base pairs) sequences of ZIKV. Hybridization of complementary sequences (colored in red) results in RNA cyclization, which is essential for RNA replication of Flavivirus. Structural elements shown include stem-loop A (SLA), 3’ stem-loop (3’ SL), start codon (AUG, colored in blue), capsid region hairpin (cHP). The two pairs of complementary sequences are 5’-3’ complementary sequences (5’-3’ CS) and 5’-3’ upstream AUG region (5’-3’ UAR). We followed previous notations used for DENV^8^. (b) Multiple sequence alignments of 5’ and 3’ terminal sequences that are involved in cyclization. The regions corresponding to 5’-3’ CS and 5’-3’ UAR are highlighted.

**Supplementary Figure 8. Consensus RNA structure of coding region near the 5’ and 3’ end**. (a) Predicted RNA structure of 300bp at the 5’ end of coding region (i.e. capsid protein). (b) Predicted RNA structure of 300bp at the 3’ end of coding region (i.e. NS5 protein). The consensus structure of strains of African lineage and Asian lineage are shown. Positions in the conserved hairpins are marked in red.

## SUPPLEMENTARY TABLE

**Supplementary Table 1**. Distribution of 3423 codons with overabundance of dN and dS and neutral codons were calculated from the different ZIKV data sets.

**Supplementary Table 2a**. Natural selection of Zika viral strains from different data sets.

**Supplementary Table 2b**. Natural selection of 11 genomic segments of Zika viral strains of Asian and West African lineages.

**Supplementary Table 3**. Details of 46 Zika viral strains studied in the present study (1947 – 2016).

**Supplementary Table 4**. Comparison of codons with excess of dN and dS from both Asian and West African lineages.

**Supplementary Table 5**. Natural selection pressure acting on each amino acid of 2015/16 epidemic Zika viral strains.

**Supplementary Table 6**. Summary of statistically reliable negatively selected amino acid sites of 2015/16 epidemic Zika viral strains. The negatively selected common amino acid sites identified by more than one method in the datamonkey were denoted by bold.

**Supplementary Table 7**. Classification of predicted CD4 and CD8 TCEs of 2015/16 BeH819966 ZIKV (KU365779) based on their immunogenicity scores.

**Supplementary Table 8a**. List of CD4 peptides predicted from 6 proteins/segments of 2015/16 ZIKV. Only immunogenic and highly immunogenic peptides with their immunogenicity scores and position in the respective protein furnished. Peptides comprise negatively selected sites were highlighted in yellow.

**Supplementary Table 8b**. List of CD8 peptides predicted from 6 proteins/segments of 2015/16 ZIKV. Only immunogenic and highly immunogenic peptides with their immunogenicity scores and position in the respective protein furnished. Peptides comprise negatively selected sites were highlighted in yellow.

---

**Supplementary Text 1: Statistical tests of positively and negatively selected sites**.

We identified 10 codons with dN>dS in both Asian and West African lineages, as well as 426 codons with dN<dS in both lineages (Supplementary Fig. 3, Supplementary Table 4). It suggests that adaptive evolutionary strategies of viruses from both Asian and African lineages were independent as they exploited only a small rational of common codons with excess of dN during the course of evolution. Notably, the codons with excess of dS exploited by two lineages are more than the codons that are unique to Asian lineage, indicate that the fraction of patterns of synonymous changes occurred in two lineage viruses are common, however the Asian strains have evolved with unique dS changes as fitness effects, because they are geographically spread in Asia and America continents.

Supplementary Table 2a shows the results of dN/dS ratio estimation. Analysis includes 46 ZIKVs (3 lineages) show that ‘ω’ ratio as 0.065, indicate clearly that ZIKVs evolved under strong purifying selection pressures. Overall results of 3 lineages show that ‘ω’ score ranged from 0.065 to 0.076, suggest that no significant differences in the overall evolution of ZIKVs belonged to different lineages. The mean of proportion of dN changes in all 3 lineages comparatively higher than the those of dS. Together, our findings showed that 2015/16 ZIKVs evolved through excess of synonymous substitutions and therefore are under strong purifying selection pressures.

The results on positive and negative selection pressures acting on each amino acid site of polyprotein encoded by 2015/16 ZIKV genomes were shown in Supplementary Tables 5 and 6. A single positively selected site (894; IFEL, *p*-value 0.08) was identified by a single method with no statistical significance. Our data showed that a total of 11 of 24 negatively selected amino acid sites were inferred with statistical significance. The present result indicates that the 2015/16 epidemic ZIKVs had evolved through purifying selection pressures, which was predicted to assist the viruses for better adaptation to the human and reduce the efficiency of active immunity. Results provided are consistent with the recent findings on Filovivirus^12^, but different from the current prevailing hypothesis that genes encoding antigens can be highly variable to evade host immunity^21-23^. In natural selection analysis, the significance of one method alone is not sufficient to infer that a given amino acid site underwent either positive or negative selection pressure^5,7^. In the present study, no positively selected sites have been detected by more than one method, whereas, out of 11 statistically reliable negatively selected sites from polyprotein, only 2 sites (360 and 3358) were identified by more than one method (FEL: *p-*values 0.05, 0.02; FUBAR: posterior probabilities 0.90, 0.98), thus, these 2 amino acid sites predicted to be more reliable. Interestingly, we were able to identify a single amino acid site (894; IFEL, *p*-value 0.08) in the polyprotein, which underwent positive Darwinian selection. Amino acid sites of ZIKV polyprotein under negative selection pressures were comparatively higher. A single negatively selected site (position 3358) identified by IFEL (*p-*value 0.07) was not statistically significant, but the same site identified by FEL (0.02) and FUBAR (posterior probability 0.98) was statistically significant. Moreover, 22 of 24 sites were identified by FEL without statistical significant. Therefore, only amino acid sites that have been detected by more than one method are finally considered as positively or negatively selected sites.

It should be noted that dN/dS tends to be biased towards one in the type of sequence samples that we had. This is because the polymorphism that we see are not actually fixed substitutions, which are assumed and generally true in the case of divergent protein sequences. This bias is independent of sample size. Another concern is the small sample size, especially for the “per site” estimation of dN and dS, which is not average over codons in the same protein. This uncertainty in the dN and dS values makes it difficult to find sites under “significant” negative or positive selection. The actual sites under negative selection can be much more.

**Estimation of relative evolutionary distance**.

In Fig. 1, the branch lengths have scaled to indicate the relative evolutionary distance between the Zika viral strains. Further comparison of genetic distance (GD)^3^ within and between the 3 lineages showed that the GD (0.011) interior of the clade Asian was differ to clade African (0.059). Within the African clade, East African and West African lineages showed 0.022 and 0.068 as GD, respectively. The GD/net GD between the Asian and East and West African clades were 0.128 (GD)/0.111 (net GD) and 0.133/0.094, respectively, whereas the GD/net GD between East and West African clades was 0.058/0.013 indicate that the ZIKVs from both African clades were genetically similar, however significant differences in their genome made them into two lineages. It is noted that the inter-lineage sequence variation in the 3 ZIKV lineages was comparatively higher than an intra-lineage sequence variations. The overall mean of base differences per site from averaging over all ZIKV sequence pairs was estimated to be 0.091 indicate that the overall sequence conservation of 46 strains studied was more.

**Supplementary Text 2: CD4 and CD8 TCEs of epidemic Zika viral isolates**.

In order to design peptide vaccine for current epidemic ZIKV and to predict the functional consequence of 11 statistically reliable amino acids under purifying selection pressure, we predicted potential CD8 (9 amino acid residues) and CD4 (15 amino acids) TCEs from 6 proteins. Our results indicated that a total of 1517 (range: 103-633) and 1481 (range: 97-627) CD8 and CD4 TCEs, respectively, were predicted. Notably, 74% (CD8 1120/1517; CD4 1096/1481) of CD8 and CD4 TCEs were predicted from the four non-structural proteins. Next, we tested the immunogenicity of these 2998 TCEs. Higher immunogenicity score indicates a greater probability of eliciting an immune response. Therefore, we grouped the peptides of immunogenicity scores <0, <0.5-0, and >=0.5 as non-immunogenic, immunogenic and highly immunogenic, respectively (Supplementary Tables 7, 8a,b). Overall results showed only a small proportion of TCEs [5 of 1517 CD8 (0.3%) and 106 of 1481 CD4 (7%)] had >=0.5 immunogenicity score.

Capsid protein contained only 3% (n=3) CD4 TCEs, whereas, envelope (central and dimerization domain) had 4% (n=12) CD4 TCEs with higher immunogenicity scores. Overall, these TCEs of C and E proteins can be utilized as vaccine candidates as most of them are located in the functional regions of the proteins. A total of 91 CD4 and 5 CD8 TCEs from 4 non-structural proteins (NS2A, NS3-serine protease, NS4A, NS5) are predicted to be highly immunogenic. Further experiments are required to validate these 111 highly immunogenic CD4 and CD8 T cell specific epitopes for induction of immune responses.

Furthermore, we evaluated the 11 amino acids that were under purifying selection pressure in the context of TCEs. These sites are less likely to generate immune-escape variants, due to strong functional constraints operating on them. 11% (n=163) and 7% (n=99) of 1481 CD4 and 1517 CD8 TCEs, respectively contain these 11 negatively selected amino acids (Supplementary Tables 8a,b). Any substitutions at these amino acid residues are likely to be lethal or intolerable^11,24,25^. For vaccine development, besides the highly immunogenic 111 TCE, additional 137 (92 CD4 TCEs; 45 CD8 TCEs) immunogenic peptides comprising negatively selected amino acid sites can be included.

**Supplementary Text 3: Caveats for interpreting site linkage in ZIKA polyprotein**.

We have identified co-evolving sites in both African and Asian genotypes. Due to the small number of sequenced ZIKA genomes, the correlation matrix is corrupted by noise after removing the dominant contributions of phylogeny (Supplementary Figs. 3 and 4). Moreover, the number of polymorphic sites in our data is small. When more sequence data are available, the dimensions of the correlation matrix will increase and the noise can be cleaned up by removing eigenvalues below the threshold given by random matrix theory^26,27^. As more ZIKA genome sequences are accumulated, we expect that these limitations will be alleviated and we will be at a better position to evaluate the influence of higher-order constraints on virus evolution (e.g. epistasis, protein-protein interactions).

Assuming that the inferred correlation between amino acid sites is real, we note the linkage in evolution can be due to reasons other than protein evolution. One interesting example is the cluster of coevolving codons 6-10 identified in the African lineage. By examining the sequence alignment, we found that ZIKA genomes had a deletion of C in codon 3 and an insertion of U in codon 11, compared to the two earliest isolated strains in 1947 (AY632535, NC_012532). The overall shift in the nucleotide sequence explains why codons 6-10 are found to be linked together in evolution. Interestingly, this pair of insertion and deletion was reverted in an Asian strain isolated in 2007 (EU545988), which is phylogenetically close to the African lineage.

Like many other RNA viruses, mutations in the coding region of ZIKV are also subject to RNA selection^28^. In the above example of deletion and insertion downstream of the start codon, we do not know whether this combination of insertion and deletion was selected due to amino acid changes of this stretch of amino acid in capsid, or due to RNA structure. RNA structures in the 5’ end coding region is important for cyclization and could potentially be interacting with nonstructural proteins in the replication complex (Supplementary Fig. 8). Thus, some linkages among amino acid sites are potentially caused by selection at the RNA level, which is a unique feature of RNA viruses.

